# Infant spatial relationships with adult males in a wild primate: males as mitigators or magnifiers of intergenerational effects of early adversity?

**DOI:** 10.1101/2024.04.25.590770

**Authors:** Matthew N Zipple, Chelsea A Southworth, Stefanie P Zipple, Elizabeth A Archie, Jenny Tung, Susan C Alberts

## Abstract

Adult male mammals can provide infants with protection and enhance their access to resources. They can also pose a risk to infants, either directly through infanticide or other aggression, or indirectly by placing infants at increased risk of conspecific or heterospecific conflict. Both benefits and costs may be especially important for offspring born to mothers in poor condition. Here we present the most detailed analysis to date of the influence of adult non-human primate males on a wide range of infant behaviors, and a description of the predictors of individual infants’ proximity to adult males. We show that the number of adult males near an infant predicts many infant behavioral traits, including aspects of the mother-infant relationship, infant activity budgets, and the frequency of social interactions with non-mothers. Infant exposure to adult males is statistically significantly repeatable over time (R = 0.16). This repeatability is partially explained by whether the infant’s mother experienced early life adversity: offspring of high-adversity mothers spent time in close proximity to more males during the first months of life. Our results are consistent with the possibility that the effects of maternal early life adversity can be mitigated or magnified by relationships with adult males.

## Introduction

Across group-living mammals, adult males pose a range of both benefits and risks to the survival of immature individuals within their groups. On the one hand, males of different species display substantial variation in the degree to which they interact with or care for immatures. Among species where males frequently encounter immatures, they often direct no recognizable care towards them (*1*, *2*). However, males of some species of group-living mammals provide extensive, even obligate, care (e.g. some mongooses (*3*, *4*), hyenas (*5*), dogs (*6*), and voles (*7*)). Paternity certainty plays a role in shaping patterns of care within and between species (*8*), but it is not determining. The distribution of paternal care is particularly well characterized in primates, which display a high degree of variation in the extent to which males care for immatures, from rare interactions in chimpanzees to direct, obligate male care in owl monkeys (*9*). Notably, even in the absence of paternity certainty, males still display care towards immatures in numerous primate species (e.g. langurs: (*10*), baboons: (*11*, *12*), macaques: (*13*), and gorillas: (*9*))

At the same time, males can pose substantial risks to infant survival. In many mammals, non-sire males kill infants to induce a return to fertility in the mothers of those infants (who are otherwise infertile during lactation) (*14*–*17*). Support for sexually selected infanticidal behavior by males is widespread across taxa, with evidence from dozens of species of mammals, including cetaceans, carnivores, tree squirrels and primates (*15*). Mothers often attempt to physically prevent such infanticidal attacks, but this protection is imperfect (*15*). Females may also associate with particular males, frequently the fathers of their infants, who serve as protectors (e.g. ground squirrels (*18*), lions (*19*), baboons (*20*), reviewed in (*15*)). As with male care, sexually selected infanticide and counterstrategies to try to prevent it have been especially well documented in primate populations (*15*, *16*, *21*).

Evaluating the relative importance of male mammals as sources of protection versus risk for infants requires both comparative data across multiple species and also detailed studies within species, which can shed light on the behavioral mechanisms that mediate benefits versus risks. Here we present such a detailed study in baboons, a species in which both the positive and negative aspects of male-immature interactions, are seen: male baboons can both provide immatures with care and protection and pose direct or indirect risks to their survival. Early in life, yellow, olive, and chacma baboon infants often consistently interact with the same male or males in their early lives, and sometimes establish individualized relationships with males (*11*, *22*–*24*). Adult males approach infants, make physical and vocal contact with them, tolerate their presence during feeding, and intervene on their behalf during agonistic interactions (*11*, *22*–*25*). Infants can substantially benefit from their relationships with males via increased access to food scraps, protection from harassment and infanticide from conspecifics, and protection from predation risk (*24*, *26*, *27*). In the Amboseli baboons of Kenya, fathers (who represent a subset of male friends) intervene on behalf of their offspring during agonistic interactions (*25*), and the presence of a father predicts earlier offspring physical maturation (*28*). Males may also provide infants with increased autonomy: infants are substantially more likely to move away from their mothers when males are nearby (*11*, *23*, *29*). However, baboon males also pose risks to infants, with sexually selected infanticide being documented in several populations (*20*, *30*). And when males engage in physical conflict with other males, they can also sometimes expose infants to incidental injury or death (*11*, *17*, *23*).

These positive and negative effects of baboon male-infant relationships may be especially important for otherwise compromised infants, such as those born to mothers in poor condition. In Amboseli, female baboons that experience early life adversity experience high adult mortality rates, leading to shorter lives (*31*). For adult females, social connections, including those to adult males, appear to reduce the effects of some forms of early adversity on longevity (*32*). Notably, the offspring of females that experience early adversity are also more likely to die during the juvenile period (*31*, *33*, *34*). The mechanisms mediating this intergenerational transmission of early adversity are unknown, but the mitigating effect of social bonds with adult males for high-adversity females suggests that males could also play an important role in either reducing or magnifying the intergenerational effect of early adversity for infants born to these mothers.

The Amboseli baboon population thus provides a unique opportunity to assess both the positive and negative aspects of male-immature interactions and the impact of the presence of adult males on immatures with different developmental histories. Here we use detailed behavioral data to makes three contributions that help us better understand the role of adult male mammals in infant development. First, we identify the ways in which adult male baboons influence infants’ lives by shaping mother-infant interactions, infant activity budgets, and social interactions between infants and conspecific groupmates. Second, we quantify the repeatability of the number of adult males in an infant’s close proximity, documenting the extent to which exposure to adult males is specific to a given infant. This is important because significant repeatability indicates that some characteristic of a mother/infant pair, such as the early adversity experienced by the mother, explains variation in infant exposure to adult males. Finally, we identify demographic and maternal factors that explain both within- and between-individual variation in infant proximity to adult males. We focus on the number of adult males in close proximity to an infant, rather than the identity of those males, as a predictor of infant behavior. This approach allows us to take a broad view of the positive and negative ways that adult males, in general, influence infant behavior and development and sets the stage for future work on relationships between infants and particular adult males.

## Methods

### Study system

The Amboseli Baboon Research Project is a longitudinal study of wild baboons living on the savanna in and around Amboseli National Park, Kenya. The study population is admixed, with both ancestral and contemporary hybridization occurring between yellow (*Papio cynocephalus*) and anubis baboons (*Papio anubis*; all individuals are admixed, with an average of 30-37% anubis ancestry (*35*)). Long-term data collection in this population began in 1971 and includes detailed demographic, behavioral, and environmental data since that time from nearly 2000 individuals (*36*). In this study, we augment the long-term data set with short-term intensive behavioral data collection to understand the causes and consequence of maternal/infant interactions with adult males.

### Study subjects and focal follow protocol

To characterize maternal-infant relationships with adult males, we performed focal follows of mother-infant pairs over 12 months, during July-October 2018, July-December 2019, and May-June 2021. We followed infants from birth or their age at the onset of the study periods up to a maximum of nine months of age. Infants were chosen by MNZ with the goal of observing infants whose mothers spanned the full spectrum of maternal rank and maternal early adversity (see below) observed in this population, distributed across infant ages. Other observers (CAS and SPC) were blind to the early adversity status of mothers.

Across the three field seasons, we performed 1213 focal follows across 66 mother-infant pairs (7-27 samples per pair, mean = 18). Of the 66 mother-infant pairs, 55 mothers are represented once, 7 mothers are represented twice, and 2 mothers are represented three times. We collected 5-19 hours of data per focal pair (mean = 12 hours), spanning 1-22 weeks of each infant’s life (mean = 9.5 weeks, Figure S1). Focal animal samples were conducted in a predetermined rotation to allow for even distribution of follows for each pair across different times of day. Most focal follows were 45 minutes long, but the final follow of the day was occasionally cut short or extended (mean focal follow length = 40.4 minutes; range = 10-58 minutes; total number of hours of focal observation). We control for differences in focal length, focal identity, and observer identity in our models, as described below.

During focal follows we collected (i) point samples, i.e., behavioral states recorded every minute, on the minute, and (ii) all-occurrence data for discrete events that were recorded each time that they happened (*37*) on a digital tablet using the Prim8 app for Android operating systems (*38*).

### Data collected as point samples

Point sample data recorded both the mother’s and infant’s instantaneous activity (resting, walking, feeding, socializing, playing) observed on the minute, for each minute in the focal sample. We also recorded whether the infant was on the nipple at the time of the point sample (yes/no), the instantaneous spatial relationship between mother and infant (in contact, separated by < 1 m, separated by 1-3 m, separated by 3-10 m, separated by > 10 m), and the number of adult males within 5 m of the infant at the time of the point sample. We defined 5 m as ‘close proximity’. When modeling the distance between mothers and infants as a response variable (see statistical methods), we assumed that infants were at the center of our distance categories (e.g. we assigned a distance of 2 m for points where infants were 1-3 m from their mothers. When infants were > 10 m from their infant, we assumed a distance of 20 m.

If a given data type could not be identified at the time of point sample, we recorded it as unknown.

### Data collected via all-occurrence sampling

During focal follows, we also recorded all occurrences of the following events: (i) each positive or negative interaction that an infant had with an individual that was not its mother, (ii) each distress call that an infant produced, (iii) each time the infant or mother made or broke contact with the other, and (iv) each time the infant or mother entered or left a 3 m radius of the other. Hereafter, we collectively refer to making or breaking contact and 3 m approaches or departures as “spatial transitions.” We use the term “approach” to mean a spatial transition in which a mother or infant reduced the distance between them and “departure” to mean a spatial transition in which that individual increased this distance.

‘Positive’ interactions occurred any time that a non-mother individual made a vocalization towards an infant or any time an infant was in contact with a non-mother individual without any distress vocalization from the infant or any attempt by the mother to prevent the interaction. Positive interactions included grooming, carrying, pulling, and playing. Negative interactions included pushing, hitting, biting, or pinning to the ground, and any of the above ‘positive’ interactions that resulted in infant distress vocalizations or maternal interference in the interaction (by pulling the infant away or turning her shoulders from the approaching individual). Interactions would sometimes begin as positive interactions (e.g., a male grunted at an infant) but then become negative interactions (e.g., the male pulled on the infant and the infant produced a distress call). In such cases, we recorded the occurrence of both a positive and negative interaction.

Positive social interactions were often multifaceted and extended, lasting more than 5 minutes in some cases. We considered an event as a distinct positive interaction, rather than a continuation of an existing interaction, if 20 consecutive seconds passed without additional positive interaction between the infant and the same individual, or if the interaction was between the infant and a new individual.

To understand the dynamics of mother-infant proximity we calculated Hinde index values using the following formula (*39*):

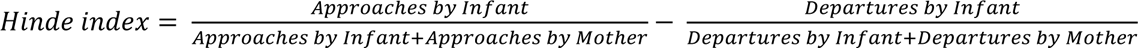

Higher Hinde index values indicate that infants are more responsible for maintaining close proximity to their mothers, while lower values indicate that mothers bear this responsibility.

### Factors that may explain variation in maternal-infant relationships with males

We tested three main types of factors that might explain variation in maternal-infant relationships with adult males: (1) maternal social dominance rank, which influences maternal resource access and rates of harassment by others, (2) maternal early life experiences, which are linked to maternal survival and social integration, and (3) social group demography, which may act as a constraint on the total number of adult males available to a given infant.

Social dominance ranks are calculated on a monthly basis for the members of each study group (see (*40*) for details). We express maternal dominance rank as the proportion of other female adults in the group that an individual dominates (*41*). This rank metric ranges from 0 to 1, with 0 representing the lowest-ranking adult female in a group and 1 representing the highest-ranking.

We measured six sources of early life adversity, calculated for mothers in each mother-infant dyad, that have previously been linked to adult female survival, reproduction, social connectedness, and/or offspring survival (*31*, *32*, *42*–*44*). Specifically, we assess whether each mother (1) experienced the death of her mother in early life (before age 4), (2) was born to a mother of a low dominance rank (proportional rank < 0.25), (3) was born to a socially isolated mother (in the bottom quartile of social connectedness), (4) was born during a drought (< 200 mm rain in her first year of life), (5) was born into a large group (top quartile of group sizes), or (6) had a close-in-age younger sibling, born within 18 months of her birth (the shortest quartile of interbirth intervals in our population). We counted the total number of adverse experiences that mothers faced to generate a cumulative measure of maternal adversity (see (*31*) for more details).

Finally, group membership is tracked on a near-daily basis by experienced observers who recognize all members of the study population on sight. We use these long-term membership data to count the number of adult males and adult females in various reproductive states that were present in a group on a given day.

### Statistical analyses

We first assessed the consequences of male presence on infant behavior by building a series of linear mixed effects models using the R package ‘glmmTMB’ (*45*). In each of 12 models, our response variable was an infant’s average behavioral measure during a focal follow. Our primary predictor variable of interest was the average number of adult males that were within 5 m of the infant (i.e., in close proximity) during the focal follow. Many of our response variables were strongly dependent on infant age, so we included infant age as a fixed effect in each model. We also controlled for the fixed effects of infant sex, maternal dominance rank, and maternal age. Finally, to separate the presence of males from the behavior of the infants’ mother, we included measures of maternal activity budget during each focal follow as an additional fixed effect.

To assess the predictors of infant exposure to adult males, we built a single linear mixed effects model (package ‘glmmTMB’) in which our response variable was the average number of adult males within 5 m of the infant in a given focal sample. To assess the differential presence of males in the lives of infants whose mothers experienced early life adversity, we included cumulative maternal early adversity and its interaction with infant age as predictor variables. We also included infant age, infant sex, maternal rank, maternal time budget, and several social demographic variables (the number of adult males, the number of pregnant and lactating females, and the number of cycling females in the social group) as fixed effects.

All linear mixed models of either infant behavior or the number of males in close proximity to an infant (i.e., within 5 m; 13 total models) included random effects of maternal identity and observer identity. We also included random slopes of infant age nested within maternal identity in those models for which adequate variance existed to support them (i.e. those models in which inclusion of a random slope did not induce a singular fit). When examination of quantile-quantile plots indicated violation of the normality assumption, we transformed response variables to improve normality of model residuals. The unit of analysis for all analyses were individual focal follows and we weighted the contribution of each follow to the log likelihood based on the number of points in the focal follow for which the response variable was known. For example, there were a total of 45,930 total points for which infant suckling status was observable across 1210 focal samples (average = 38 minutes per sample). A focal sample that included 38 minutes of known suckling status was assigned a weight of 1, while a sample that included only 19 minutes of known suckling status was assigned a weight of 0.5, using the ‘weights’ argument within the glmmTMB function.

Finally, to assess the extent to which the average number of adult males within 5 m of a particular mother-infant pair was a repeatable characteristic of that pair, we calculated repeatability using the R package ‘rptR’ [53]. When calculating repeatability we included infant age as a fixed effect.

## Results

### I. The influence of males on infant behavior and social development

In the aggregate, infants of all ages spent 23% of their time in proximity (i.e., within 5 m) to 1 or more adult males (1 male: 19%, 2+ males: 4%). However, infant proximity to males depended on infant age: infants spent less time near adult males as they aged (see below for more details), declining from a high of 29% of their time in their first 3 months of life (2+ males: 5.3%) to 19% during months 6-9 (2+ males: 2.7%). See Table 1 for additional summary statistics.

**Table 1.** Descriptive statistics of the proportion of time that infants of different ages spent within 5 m of different numbers of adult males.

| Infant Age (months) | Time Spent Within 5 m of Adult Males<br>% of Point Samples (# of Point Samples) |  |  | Mean Number of Adult Males Within 5 m |
| --- | --- | --- | --- | --- |
|  | 0 Males | 1 Male | 2+ Males |  |
| 0.0-2.9 | 71% (13412) | 24% (4495) | 5.3% (1009) | 0.35 |
| 3.0-5.9 | 80% (7639) | 17% (1570) | 3.1% (294) | 0.23 |
| 6.0-8.9 | 81% (14086) | 16% (2758) | 2.8% (484) | 0.22 |

The behaviors of individual infants differed when they were in close proximity to relatively many males versus when they were not near adult males (Figure 1). When infants were in proximity to many adult males they spent less time on their mother’s nipple (p < 0.0001, Figure 1A, Tables 1, S1), were farther away from their mother (p < 0.0001, Figure 1B, Tables 1, S2), were less responsible for maintaining contact with their mothers (p = 0.03, Figure 1C, Tables 1, S3), and spent more time engaged in active behaviors (i.e., not resting, p < 0.0001, Fig. 1D, Tables 1, S4). This increase in active behaviors included increases in time spent feeding (p = 0.01), walking (p = 0.0002), socializing (p < 0.0001), and playing (p < 0.0001, see Table S4 for additional details).

**Figure 1.**
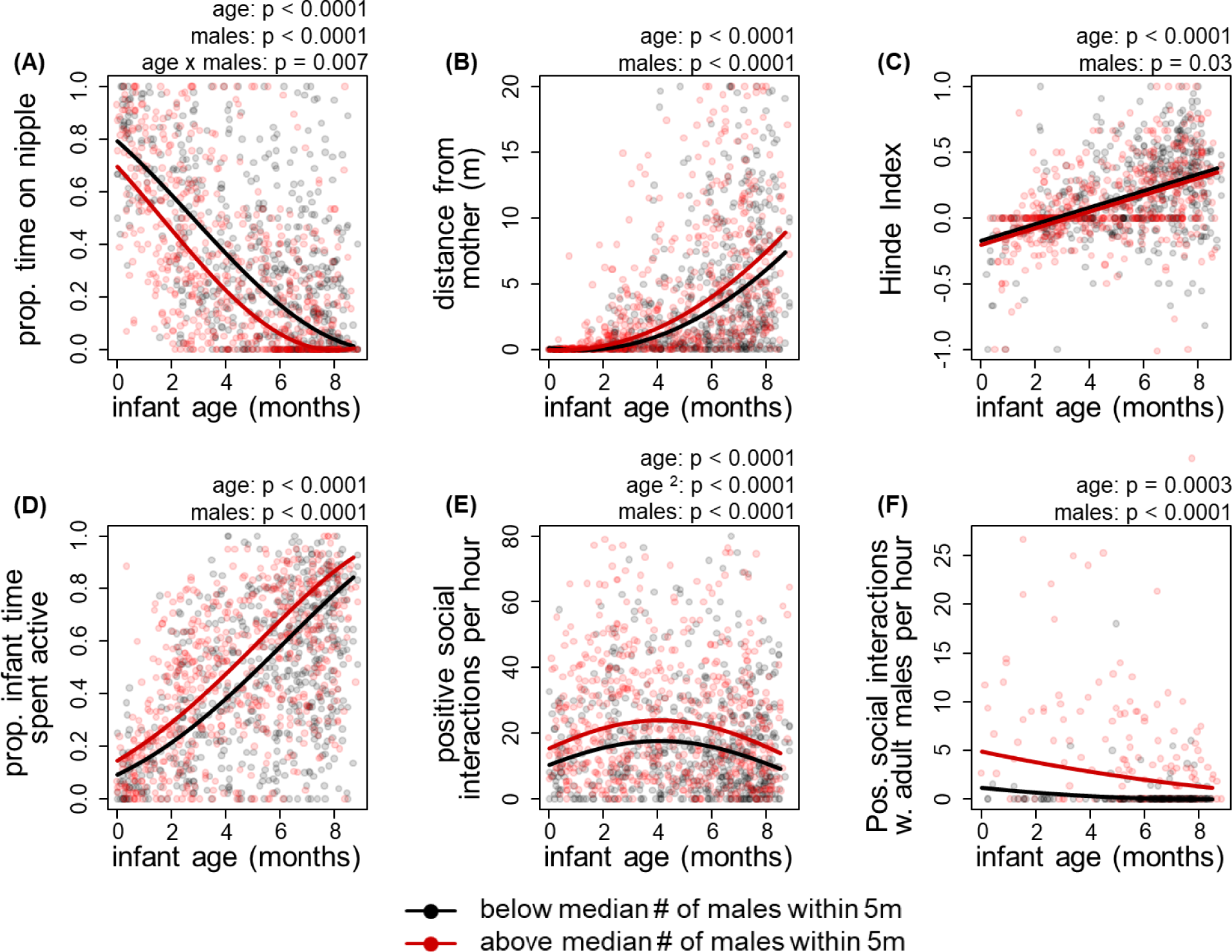
Consequences of male presence for infant behavior. When infants were in close proximity to more males, (A) infants spent less time on the nipple, (B) infants were further from their mothers, (C) mothers were more responsible for maintaining proximity to infants, (D) infants spent more of their time active, (E) infants were engaged more frequently in positive social interactions with non-mothers, and (F) infants were specifically engaged more frequently in positive social interactions with adult males. In all panels, the p values indicate the results of the full models reported in Tables 1, S1-S6. Points in each panel represent average values for a given focal follow. For visualization purposes, curves indicate estimates from simplified models (not the full models reported in the Supplemental Tables), which include only infant age, the average number of males within 5m of the infant, and their interaction (if applicable), along with random effects of mother ID and a random slope of infant age (if applicable). For visualization purposes, y-axes are truncated in (E) and (F), causing us to exclude 2% and 5% of data points, respectively. These values were retained the statistical analyses.

Interpreting the magnitude of the association between male presence and infant behaviors is challenging because some of our response variables were transformed, all models had multiple covariates, and infant behaviors were age-dependent. We therefore sought to generate a heuristic to assist in interpretation. For age-dependent behaviors, we compared the coefficient estimates from our models of the effect of adult males on infant behavior to the coefficient estimates of infant age from the same model. This approach yields an estimate of how much older or younger an infant ‘appeared’ when they were in the presence of adult males. Although these interpretations do not take into account uncertainty in the coefficient estimates, they provide a coarse, but informative, tool for interpreting the magnitude of the differences in infant behavior in the presence of adult males.

For example, at birth, an infant who was in proximity to 1 adult male was estimated to spend the same amount of time on the nipple as that same infant when it was 2.7 months older but in proximity to zero adult males (magnitude of 1 adult male = -0.30, magnitude of 1 month = -0.11). The magnitude of this effect declined as infants get older (infant age x males p = 0.007, Table 1), such that a 6-month-old infant in proximity to 1 adult male was on the nipple as frequently as when that same infant was 0.9 months older but in proximity to zero adult males. Similarly, an infant exposed to 1 adult male was estimated to be the same distance from its mother as when that same infant was 1.6 months older but in proximity to zero adult males. Such an infant spent as much time active as that same infant when it was in proximity to zero adult males but was 1.8 months older, including increased time feeding (0.8 months older), walking (1.5 months older), and playing (3.6 months older). In contrast, an infant in proximity to one adult male was less responsible for maintaining contact with its mothers (i.e. such a pair showed a lower Hinde index) when an additional male was present, comparable to the same infant that was 1.0 months younger but in proximity to zero adult males.

Proximity to adult males also predicted differences in infant social behavior. Infants spent more time engaged in non-play social interactions when exposed to more adult males (p < 0.0001, estimated average time socializing = 4.6%; for infants in proximity to one additional male than average infant = 8.0%). In absolute terms, infants were involved in significantly more social interactions, both positive and negative, during focal follows when they were in close proximity to more adult males (p < 0.0001, Figure 1E, Figure S2, Tables 1, S5). The average infant is predicted to have engaged in 31% more positive interactions per hour (5.45 vs 4.17) and 67% more negative interactions per hour (0.72 vs 0.43) for each additional adult male in its proximity (note that negative interactions, at fewer than one per hour on average, were rare compared to positive interactions, which occurred 4-5 times per hour on average)

The increase in social interactions when infants were in close proximity to more males was primarily driven by an increase in interactions with males. For the subset of follows where the age class and sex of the interactant was known (n = 296 follows from the data collected in 2019), we found that infants engaged in more positive social interactions with adult males when they were in proximity to more adult males (p < 0.0001, Figure 1F, 350% increase in such interactions for each additional male present). However, they did not engage in more positive social interactions with other individuals (p > 0.1, Tables 1 and S6). We did not collect data on the age-sex class of interactants involved in negative social interactions.

Finally, we tested whether the presence of adult female neighbors (not including the mother) had similar effects as adult male neighbors, using the subset of follows in 2021 for which we also recorded the number of adult females within 5 m of the focal infant (n =289). We found that proximity to adult females had some, but not all, of the same effects on infant behavior as did adult males. Like males, proximity to adult females predicted lower mother-infant Hinde Index values (p < 0.05), more infant time spent playing (p < 0.001), and an increased number of positive social interactions (p < 0.0001). On the other hand, the number of adult females in close proximity to infants did not predict the amount of time that infants were on the nipple, time they spent active, or their distance from their mother (p > 0.05 for each). Overall, our results demonstrate that while proximity to adult males and proximity to adult females show some overlap in the ways that they predict infant behavior, adult males play a unique role in infant development (Table 3, full results: Table S7).

**Table 2.** Summary of effects, on infant behavior, of the number of adult males within 5m of the infant. Each row corresponds to the effect of number of adult males on one of eight different measures of infant behavior, in one of eight statistical models. In this summary table we report the coefficient ± standard error and the t-value and p-value (bolded for < 0.05) for the effect of number of adult males within 5 m. Note that in one model the effect of an interaction between infant age and number of adult males was statistically significant; this result is presented in a separate column. No other model produced statistically significant interactions with infant age. In each model, maternal time resting, walking and socializing were included as controls to allow us to differentiate effects of maternal activity from effects of adult males. Full models for each response variable are presented in Supplemental Tables S1-S6.

| Response variable | Transformation of response variable | No. of adult males within 5m of subject | Interaction: [infant age] x [no. of adult males within 5m] | Interpretation | Full model presented |
| --- | --- | --- | --- | --- | --- |
| Proportion of time that infants spent on the nipple | Arcsine square root transformed | Coef. = $-0.30 \pm 0.06$<br>t = -5.91<br>**p < 0.0001** | Coef. = $0.03 \pm 0.01$<br>t = 2.68<br>**p = 0.007** | Infants spent less time on the nipple when in proximity to more adult males. Newborns in proximity to one additional adult male behaved as if they were 2.6 months older. 6-month-olds behaved as if they were 1.0 month older. | Table S1 |
| Distance between an infant and its mother | Square root transformed | $0.58 \pm 0.08$<br>6.85<br>**p < 0.0001** | N/A | Infants were farther from their mothers when in proximity to more adult males. Infants in proximity to one additional adult male behaved as if they were 1.6 months older. | Table S2 |
| Hinde Index of mother-infant proximity | Untransformed | $-0.06 \pm 0.03$<br>-2.20<br>**p = 0.03** | N/A | Mothers were more responsible for maintaining proximity when more males were present near the infant. When one additional male was present, mothers behaved as if their infants were 1.0 months younger. | Table S3 |
| Proportion of time infants spent active | Arcsine square root transformed | $0.15 \pm 0.02$<br>5.97<br>**p < 0.0001** | N/A | Infants spent more time active when in proximity to more adult males. Infants in proximity to one additional adult male behave as if they were 1.8 months older. | Table S4 |
| Frequency of positive social interactions with non-mothers | Square Root Transformed | $1.29 \pm 0.19$<br>6.88<br>**p < 0.0001** | N/A | Infants more frequently engaged in positive social interactions when in proximity to more adult males. The average infant is estimated to have engaged in 31% more positive interactions per hour (5.45 vs 4.17) if in proximity to one additional male. | Table S5 |
| Frequency of negative social interactions with non-mothers | Square Root Transformed | $0.68 \pm 0.15$<br>4.54<br>**p < 0.0001** | N/A | Infants more frequently engaged in negative social interactions when in proximity to more adult males. The average infant is estimated to have engaged in 67% more negative interactions per hour (0.72 vs 0.43) if in proximity to an additional male. | Table S5 |
| Frequency of social interactions with adult males | Square root transformed | $3.2 \pm 0.24$<br>13.61<br>**p < 0.0001** | N/A | Infants more frequently engaged in social interactions with adult males when in proximity to more adult males. The average infant is estimated to have engaged in 350% more positive interactions with males per hour (4.13 vs 0.91) if in proximity to an additional male. | Table S6 |
| Frequency of social interactions with non-mother females and immatures | Square root transformed | $0.78 \pm 0.52$<br>1.48<br>p = 0.14 | N/A | No effect | Table S6 |

**Table 3.** A comparison of the effect that the number of adult males and adult females have on infants (for those effects of males that could be reproduced in the subset of follows containing data on the number of adult females).

| Measured Outcome | Effect of Adult Males | Effect of Adult Females |
| --- | --- | --- |
| Time on Nipple | ↓ | None |
| Distance From Mother | ↑ | None |
| Hinde Index* | ↓ | ↓ |
| Time Spent Active | ↑ | None |
| Time Spent Feeding | ↑ | None |
| Time Spent Playing | ↑ | ↑ |
| Positive Social Interactions Per Hour | ↑ | ↑ |

\*The model using only the subset of follows that contained data on the number of females within 5 meters of the infant *did not* reproduce the effect of adult males apparent in the full dataset. Despite this reduced statistical power, the number of adult females in close proximity still predicted the Hinde Index, so we include this result here.

### II. Determinants of infant proximity to males

#### (a) Mother-infant pair identity

The mean number of adult males within 5 m of an infant was statistically significantly repeatable within individual mother-infant pairs (p < 0.0001), indicating that one or more specific characteristics of the mother/infant pair are predictive of infants’ exposure to adult males. After controlling for infant age and observer identity, mother/infant identity explained approximately 16% of the total variance in the number of males within 5 m of an infant during a focal follow (R = 0.16, 95% CI = [0.10, 0.22]). Figure 2 shows examples of infants that were consistently in close proximity to relatively many or few adult males during their multiple focal follows. The demographic composition of social groups explained some of this variance (see below), but mother/infant identity continued to explain 8% of total variance after controlling for the number of adult males and pregnant or lactating females in each group (p < 0.0001, R = 0.08, 95% CI = [0.04, 0.13).

**Figure 2.**
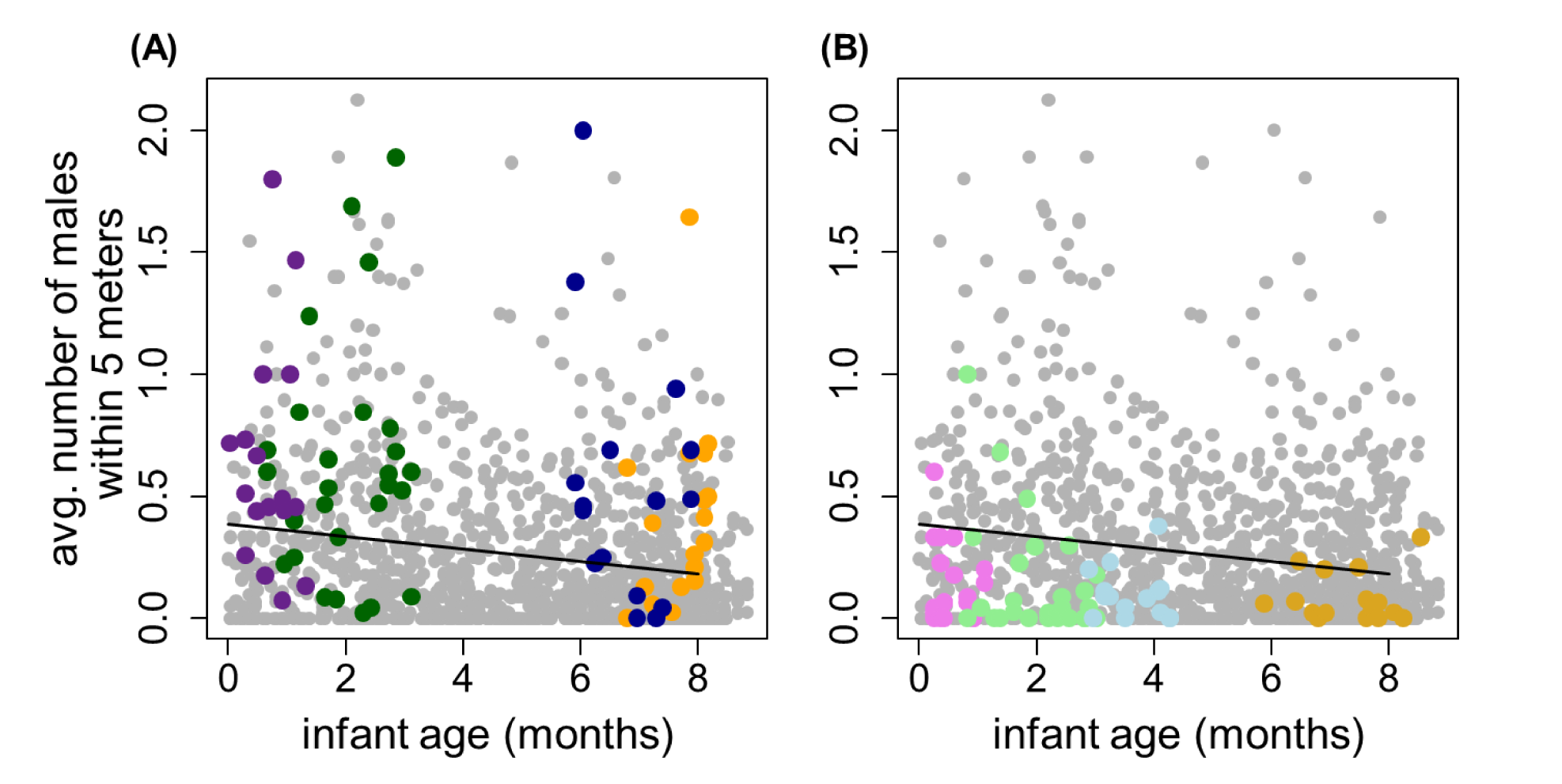
The number of males that an infant is in close proximity to is significantly repeatable (R = 0.16). Panel (A) shows four example mother-infant pairs for infants that were consistently in close proximity to relatively many males (bright colored points), while panel (B) shows four example mother-infant pairs for infants that were consistently in close proximity to few males (pastel-colored points). Gray points represent all focal follows of all mother-infant pairs in the dataset (n=1213 follows, 66 mother-infant pairs pairs) and the black line indicates the age-specific population mean.

#### (b) Social group demographics

The demographic characteristics of the social group strongly predicted the number of adult males within 5 m of an infant during focal follows (Table 4). As the number of adult males in a social group increased, so did the number of males within 5 m of the focal infants (coefficient estimate = 0.028 for each additional male in the group, p < 0.0001). However, as the number of pregnant and lactating females in a group increased (holding the number of adult males constant), the number of males to whom an infant was in close proximity declined (coefficient estimates = -0.014 for each additional female, p < 0.0001). The number of cycling females in the group did not predict infants’ proximity to adult males. The result of these combined demographic effects is that membership in a larger social group did not reliably predict increased proximity between infants and adult males. Rather, infants were in close proximity to relatively many males when the number of pregnant and lactating females relative to the number of adult males (which we term the number of ‘excess’ pregnant and lactating females) was low (see Figure 3).

**Figure 3.**
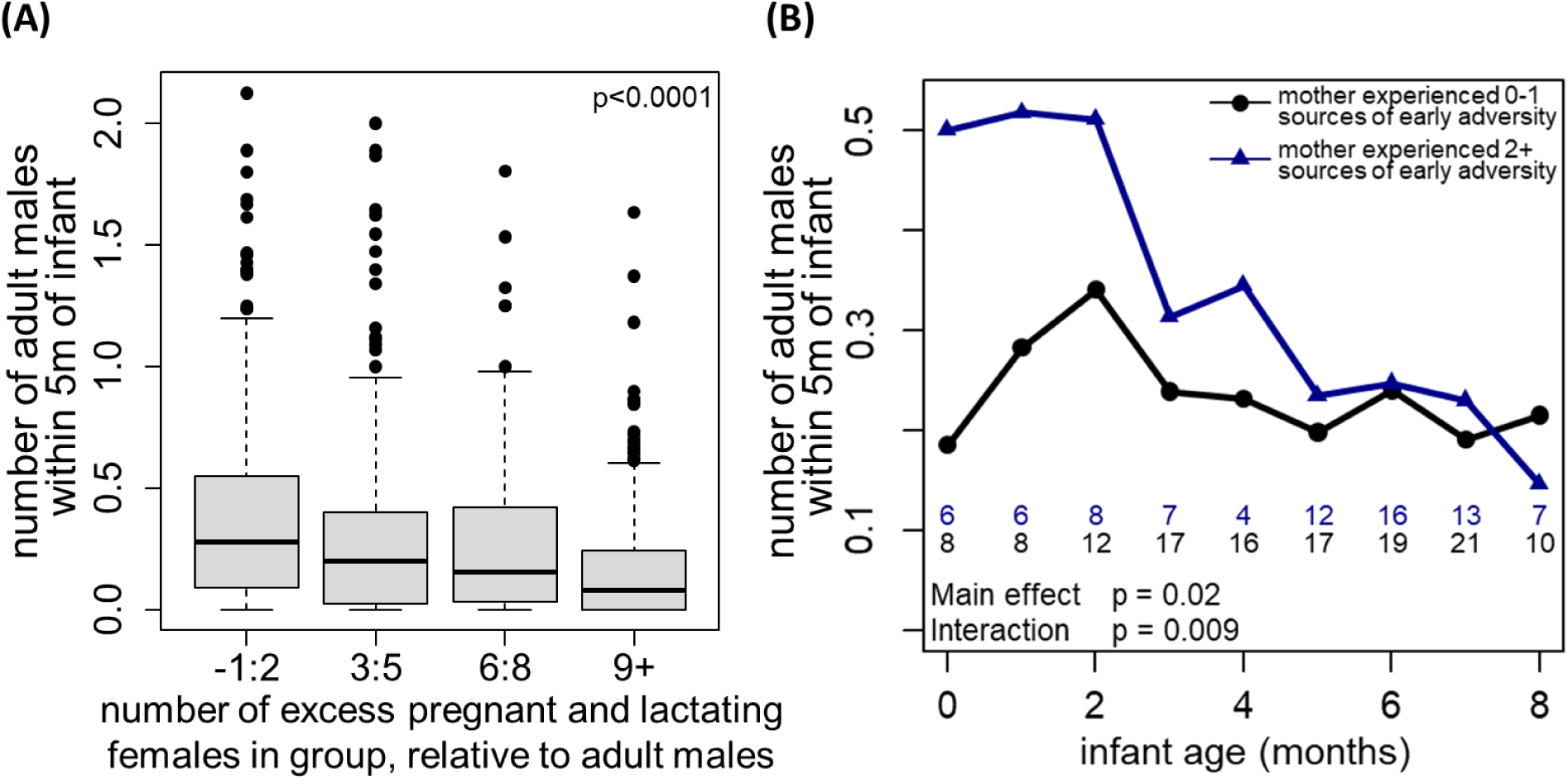
Predictors of male presence in close proximity to infants. **(A**) The number of adult males in close proximity to the focal infant decreases as the number of ‘excess’ pregnant and lactating females in a social group increases (i.e., number of pregnant+lactating females per adult male). **(B)** Cumulative maternal early life adversity strongly predicts the number of males in close proximity to infants during early life, an effect that dissipates as infants get older. Each point shows the average number of males across all focal follows where infants’ ages fell within a given one-month window (e.g., when x = 0 infants are 0 and 1 month of age). Blue and black numbers below the curve refer to the number of infants represented in each point. Although binned in these figures, predictor variables are treated continuously in our statistical analyses, reported in Table 4.

**Table 4.** Predictors of the average number of males within 5 meters of an infant during a focal follow (square root transformed).

| Parameter<sup>^</sup> | Coefficient Estimate<sup>*</sup> | Std. Error | t value | p value | Interpretation |
| --- | --- | --- | --- | --- | --- |
| Intercept | 0.35 |  |  |  |  |
| ***Social Group Demography*** |  |  |  |  |  |
| Number of Adult Males | 0.027 | 0.006 | 4.47 | **<0.0001** | Infants were in proximity to more males when there were more adult males in their social group. |
| Number of Pregnant and Lactating Females | -0.014 | 0.003 | -4.32 | **<0.0001** | Infants were in proximity to fewer males when there were more pregnant and lactating females in their social group. The magnitude of this effect is approximately half that of an additional adult male, suggesting that a given male is unable to divide his attention between more than two pregnant and lactating females at a time. |
| Number of Cycling Females | 0.004 | 0.006 | 0.70 | 0.48 |  |
| ***Maternal and Infant Characteristics*** |  |  |  |  |  |
| Infant Age (Months) | 0.0003 | 0.009 | 0.04 | 0.97 |  |
| Infant Sex (Female Reference) | 0.02 | 0.02 | 0.63 | 0.53 |  |
| Maternal Age (Years) | -0.0002 | 0.003 | -0.07 | 0.94 |  |
| Maternal Dominance Rank | 0.04 | 0.05 | 0.78 | 0.44 |  |
| Cumulative Maternal Early Adversity | 0.08 | 0.03 | 2.42 | **0.02** | Infants born to high-adversity mothers were in proximity to more males during the earliest months of life. |
| Cumulative Maternal Early Adversity x Infant Age | -0.014 | 0.005 | -2.62 | **0.009** | The difference in the number adult males in proximity to infants born to high-adversity mothers declined as infants age, such that the effect was estimated to be zero by six months of age. |
| ***Maternal Time Budget Controls (Reference = 100% feeding)*** |  |  |  |  |  |
| Prop. Time Mother Spent Resting | 0.07 | 0.05 | 1.32 | 0.19 |  |
| Prop. Time Mother Spent Walking | -0.12 | 0.05 | -2.49 | **0.02** | Infants were in proximity to fewer males when their mothers were walking |
| Prop. Time Mother Spent Socializing | 0.15 | 0.06 | 2.30 | **0.02** | Infants were in proximity to more males when their mothers were socializing. |

<sup>^</sup>Model also includes random effects of mother ID with a random slope of infant age as well as observer ID
<sup>\*</sup>Observations are weighted by the number of minutes in each focal follow where the number of adult males in proximity to the infant could be observed

#### (c) Maternal early life experience and infant age

Cumulative maternal early adversity was a statistically significant predictor of the number of adult males within 5 m of an infant. Specifically, newborn infants were in proximity to more males if their mothers experienced greater cumulative early adversity (coefficient estimate = 0.08, p = 0.02). This effect declined significantly with infant age (Figure 3B, interaction coefficient estimate = -0.014/month, p = 0.009), such that infants were no longer differentiated based on maternal early adversity by age six months. Infant age was not a significant predictor of close proximity to males independent of this interaction (e.g., among infants whose mothers experienced zero sources of early adversity; p = 0.97).

To identify the specific sources of adversity that drove the cumulative adversity-infant age interaction, we conducted a *post hoc* multivariate analysis that treated each source of early adversity separately. We found that newborn infants were in proximity to more males if their mothers experienced drought, maternal loss, or social isolation early in life, and in proximity to fewer males if their mothers were low status early in life (p < 0.05, details in Table S8). Like our results for cumulative adversity, the maternal drought effect attenuated with age (p < 0.05), whereas for the other three sources of maternal early adversity (maternal loss, social isolation, social status), we identified no significant interactions with age (p > 0.05).

## Discussion

We have shown that the number of adult male baboons in close proximity to an infant strongly predicts a range of fundamental aspects of infant early life behavior, including an infant’s activity budget, its relationship with its mother, and its interactions with non-mother individuals (summarized in Table 2). While proximity to adult females is similarly associated with some of these infant outcomes, it does not predict all of them, indicating that adult males may play a unique role in the behavioral development of infant baboons (Table 3). The relationship between infant behavior and infant proximity to males could result from a pattern in which adult males are simply more attracted to active infants, rather than male presence stimulating infant activity. While we cannot entirely rule out this possibility, the structure of our model controls for a random effect of maternal identity, which in most cases is also equivalent to the maternal-infant pair since most mothers are represented once. Consequently, our results largely reflect how male presence predicts differences in activity within a given infant, suggesting that it is male presence that drives differences in behavior, rather than stable infant characteristics that drive activity levels and therefore attraction of males.

Furthermore, infant baboons showed consistent inter-individual differences in the extent to which they were in close proximity to adult males. These differences between infants are partially explained by social group demographics, specifically the number of adults of both sexes in the group. They are also partially explained by the early life experiences of their mothers (Table 4). Infants whose mothers experienced higher levels of early-life adversity were in close proximity to more males in the earliest months of life, a difference that declined with infant age. We discuss each of these results in more detail below.

### Effects of adult males on offspring behavior and experience

One interpretation of our results is that the protection that adult males provide can ‘activate’ infants and allow them the freedom to explore and interact with their environment (*11*, *23*, *47*). Infant baboons spend less time resting and more time walking, feeding, socializing, and playing when they are in close proximity to more males, and they do so at a greater distance from their mothers. While some of the increase in social behavior and positive social interactions is linked to direct interactions with adult males, the increase in non-social active behaviors indicates that males’ influence extends well beyond direct interactions with infants. For example, the increase in play behavior that accompanies the presence of greater numbers of males is explicitly *not* the result of direct interaction with males, but instead indicates increased interactions with other infants and juveniles (Table S4). This result is similar to findings from anubis baboons, in which infants with less responsive mothers are more likely to develop ‘secondary attachments’ to non-mother social partners and display increased play behavior (*48*).

This putative activating effect of adult males has been described in human development (termed ‘the father-child activation relationship’ (*49*–*51*)). Fathers’ encouragement of children to take risks in humans is assumed to be balanced against a concern for child safety and protection (*49*–*51*). Given paternity uncertainty, baboons’ reduced theory of mind as compared to humans (*52*) and the increased dangers of the open savannah as compared to typical modern human environments, the motivation and ability of males to protect the immatures that they interact with is likely to be dramatically diminished in our study system relative to human societies. As a result, the activating effect of males that we observe may be largely beneficial for infants, but infants may also be more frequently exposed to high-variance, unpredictable scenarios at a younger age than they might be in the absence of adult males.

For example, a young infant who has strayed far from its mother in the presence of a male might find itself unable to return to its mother quickly enough if it experiences a sudden attempted predation event. Similarly, such an infant may become involved, either directly or indirectly, in an aggressive encounter between two males (reviewed in (*17*)). These encounters can be quite violent, even resulting in infant death (*53*), and a young infant might find itself unprepared to navigate such an extreme and fast-paced social encounter. In addition, achieving greater independence at a younger age – including spending less time on the nipple – may result in other physical vulnerabilities, including marginally poorer nutrition or hydration, or smaller body size (*54*, *55*). While these effects may be insignificant on any given day, they may place infants in a more vulnerable state, for instance on days when the social group fails to visit a water hole, or when a rainy season fails suddenly. Our results consistently show that infants behave as if they are older when they are in the presence of adult males, and this change in behavior may come with the risk of facing experiences and situations that an infant is not yet capable of handling safely.

Some changes in maternal behavior in the presence of adult males are likely involved in mediating the activating effect of males. Especially in the first months of life, mothers can physically prevent their infants from leaving their side, suggesting that maternal permission is a prerequisite for infants’ increased autonomy. Mothers do not always allow such increased autonomy in the presence of adult males: when more males are in close proximity to an infant, mother-infant pairs show lower Hinde index values, indicating that mothers are more responsible for maintaining close proximity to their infants (Figure 1C, Table 2). Vervet monkey mothers show a similar pattern following the arrival of a new male to their social group, increasing the frequency with which they make contact with young infants when such males are present (*56*)

Mothers’ increased protectiveness, at least in response to some males, appears warranted: increased male presence is correlated with an increase in some negative outcomes for infants in this and other nonhuman primate populations (*11*, *17*, *23*, *53*, *56*). In our study, follows in which infants were in close proximity to relatively many adult males involved more frequent, and sometimes dramatic, negative social interactions (Table 2, Figure S2). On two occasions, we observed adult males ‘kidnapping’ focal infants—events in which males carried infants from their mothers and prevented both mother and infant from reuniting with each other for an extended period of time. These episodes were protracted (in one case a male appeared to have held an infant overnight), were highly distressing to both infants and mothers, and presented an acute mortality risk to infants in the form of starvation or dehydration; detailed descriptions of these events are in the Supplement. Indeed, at least one such kidnapping by an adult male, in 2013, resulted in the infant’s death, and two other such events were associated with infant death within two days of the kidnapping (unpublished data of the Amboseli Baboon Research Project). More generally, the ability of mothers to protect their infants from infanticide by newly immigrant males depends partially on their ability to keep distant from those males and the presence of fathers that provide protection from them (reviewed in (*17*)).

### Determinants of infant proximity to adult males

#### Variation among infants

Some infants were consistently in close proximity to more males while others were consistently in proximity to fewer males (R = 0.16). About half of this repeatability was attributable to characteristics of the focal social group. In groups with a relatively high number of pregnant and lactating females relative to the number of males in the group (i.e., a higher number of ‘excess’ females), infants were more rarely in close proximity to adult males. Male baboons may themselves experience fitness benefit by providing direct and indirect care to immatures, whether or not those immatures are their offspring (*11*, *22*–*25*)); they may therefore benefit from spreading their attention to multiple infants simultaneously. If so, the pattern we describe here suggests that males are constrained in their ability to do so. However, the number of males in close proximity to an infant remained significantly repeatable after controlling for these demographic characteristics of the social group (R = 0.08). Given the strong and multifaceted downstream effects of infant proximity to males, these individual differences in infant exposure to males may set offspring on differential social or physical developmental trajectories, with long-term consequences.

#### Effects of maternal early adversity

As noted earlier, previous research in the study population indicates that infants whose mothers experienced early life adversity face elevated mortality during the juvenile period relative to other infants (*33*). The results we present here indicate that young infants also spent time in proximity to more males if their mothers experienced higher levels of early adversity. Taken together, these results are consistent with the possibility that increased exposure to adult males in early life plays a role in offspring survival in this population. Specifically, our results support the idea that adult males may either mitigate or magnify the disadvantages faced by infants born to mothers who experienced high levels of early life adversity. More extensive collection of behavioral data, combined with lifetime survival data for the infants studied here will shed additional light on whether proximity to males is ultimately a mitigator or magnifier of disadvantages associated with having a high-adversity mother is an important topic for future study.

#### Male presence as low-cost male care

Our results are consistent with the hypothesis that males shape infant behavior and development simply by being physically present in close proximity to an infant (often referred to as “indirect care” (*57*, *58*)). The protection that males provide infants seems to be largely passive, and is therefore likely to be low-cost, or even cost-free, except for the opportunity cost that males pay by not actively engaging in reproductive effort at the time (*29*, *57*–*59*). Our results provide a detailed description of how even indirect effects of male presence can influence immatures, a subject that is under-studied as compared to more overt, direct care (see (*9*, *60*)).

Because of the design of our behavioral protocol, we were not able to identify levels of genetic relatedness between focal infants and adult males with whom they were in close proximity. However, previous work has shown that infants’ adult male friends are often – but not always—their fathers (*12*, *27*, *61*, *62*). Our results therefore add to the growing body of evidence from both this population and other cercopithecines that paternal care, especially low-cost paternal care, is not restricted to socially monogamous species (*25*, *27*, *28*). The depth and taxonomic breadth of such paternal care is likely greater than is currently realized (*57*).

## Acknowledgments

Short-term data collection was supported by an NSF DDRIG (grant # 1826215), and a student grant from the Leakey Foundation. We are also grateful for the support of a pilot grant from the Duke Population Research Center (DPRC) and its NICHD Center Grant (P2C HD0065563). MNZ was supported by an NSF Graduate Research Fellowship, a Klarman Postdoctoral fellowship at Cornell University, and an NSF Postdoctoral Research Fellowship in Biology (grant # 2109636). We gratefully acknowledge the support of the National Science Foundation and the National Institutes of Health for the majority of the data represented here, currently through R01AG053308, R01AG071684, R01AG075914, and R61AG078470. Current support for field-based data collection also comes from the Max Planck Institute for Evolutionary Anthropology, and we thank Duke University, Princeton University, and the University of Notre Dame for financial and logistical support. In Kenya, our research was approved by the Wildlife Research Training Institute (WRTI), Kenya Wildlife Service (KWS), and the National Commission for Science, Technology, and Innovation (NACOSTI). We also thank the University of Nairobi, the Institute of Primate Research (IPR), the National Museums of Kenya, the members of the Amboseli-Longido pastoralist communities, the Enduimet Wildlife Management Area, Ker & Downey Safaris, Air Kenya, and Safarilink for their cooperation and assistance in the field. Particular thanks go to the Amboseli Baboon Project long-term field team (R.S. Mututua, S. Sayialel, J.K. Warutere, I.L. Siodi, I.L., and L. Musembei), and to T. Wango and V. Oudu for their untiring assistance in Nairobi. The baboon project database, Babase, is expertly managed by N. Learn, J. Gordon, and W. Wilbur. Database design and programming are provided by K. Pinc. This research was approved by the IACUC at Duke University, University of Notre Dame, and Princeton University and the Ethics Council of the Max Planck Society and adhered to all the laws and guidelines of Kenya. For a complete set of acknowledgments of funding sources, logistical assistance, and data collection and management, please visit http://amboselibaboons.nd.edu/acknowledgements/.

## Data Availability

During peer review, Anonymized data and code supporting the analyses in this manuscript is available at: https://cornell.box.com/s/6sdgw8gmal5rzal4fm8j2zwg7i0blsth

## Supplementary Materials, containing Supplementary Text, Supplemental Tables S1-S8 and Supplemental Figures S1-S2

### Supplementary text

#### Description of observed infant kidnapping behavior by males

On two occasions, adult males kidnapped focal infants during follows. We define kidnapping here as an event lasting greater than five minutes in which an individual maintained contact or close proximity with an infant while preventing the mother of the infant from making contact with the infant. Such events are typically accompanied by repeated distress calls from the kidnapped infant and may also involve attempts by the mother to reclaim the infant.

The first kidnapping occurred on August 1, 2018 and began as an extended grooming interaction between an adult male (VEB) and the focal infant (UNC), who was 11 days old at the time. VEB had been previously observed committing one of the few infanticidal attacks ever directly observed in our population (most infanticidal attacks are unobserved; Zipple et al. 2017). Specifically, in April 2012, VEB immigrated into a new social group and was observed killing and eating APR, a 3-month-old infant that he had not sired, in October 2012, six months after he immigrated. As in the case of infant APR, VEB was unlikely to have been the father of UNC: although he was living in the group at the time of UNC’s conception, he was not observed to be a consort partner of UNC’s mother, LUX, during the estrus cycle in which UNC was conceived.

When UNC attempted to break contact with VEB and make contact with his mother (LUX), VEB prevented UNC from doing so. Despite attempts by LUX to reclaim UNC from VEB, the kidnapping continued for more than one hour (beyond the period of the focal follow). During this time UNC gave frequent distress calls. UNC was unable to suckle during this time, and UNC was repeatedly placed at risk of substantial injury. UNC was kidnapped by VEB on at least two more occasions over the next 10 days and in each case was unable to obtain milk during the duration of the kidnapping events. These other two kidnappings did not occur during focal follows of UNC, so details of these events are more limited. The duration of these other events is unclear, but one or both likely included separation from LUX overnight (> 12 hours without milk or contact with mother). UNC survived infancy and was still alive at age five years when last observed.

The second kidnapping that occurred during a focal follow occurred on September 12, 2018 and involved a 24-day-old infant (VOZ), who was separated from her mother (VEJ) by a substantial distance for approximately 30 minutes. Unlike in the kidnapping of UNC, the male involved (whose identity was unknown to the observer at the time) released VOZ and allowed her mother to reclaim her. VOZ survived to 14 months of age, when she was fatally injured in an event not witnessed by observers; circumstantial evidence points to an immigrant male who was seen attacking other infants around the time of VOZ’s death.

These two kidnapping events had a clear behavioral signature in our dataset. Focal follows of infants that were less than a month old and involved a kidnapping event featured infants that were much further from their mothers on average (0.77 m and 5.74 m) as compared to all other follows of similarly aged infants (mean = 0.01 m, SD = 0.03 m, median = 0 m, max = 0.11 m). Focal follows involving a kidnapping also showed much lower proportions of infant time spent on the nipple (0.07 and 0.40) as compared to follows of similarly aged infants (mean = 0.79, SD = 0.17, median = 0.84, min = 0.32). Finally, kidnapping events resulted in infants producing many more distress calls per hour (34.1, 6.7) as compared to follows of similarly aged infants (mean = 2.2, sd = 4.5, median = 0, max = 21.6).

While it is not clear whether these specific kidnapping events had any negative long-term consequences for the infants involved, kidnapping by males and females is an acutely dangerous event for young infants, who are at risk of starvation and dehydration without access to their mother’s milk, and fatal kidnappings have been previously reported in this population (*63*, *64*).

### Supplementary Tables

**Table S1.** Parameters from a mixed effects linear model of the proportion of time that infants spend on the nipple during a focal follow (arcsine square root transformed).

| Parameter<sup>^</sup> | Coefficient Estimate* | Std. Error | t value | p value |
| --- | --- | --- | --- | --- |
| Intercept | 0.98 |  |  |  |
| ***Predictor Variables of interest*** |  |  |  |  |
| Infant Age (Months) | -0.11 | 0.009 | -13.4 | **<0.0001** |
| Infant Sex<br>(Reference = Female) | 0.03 | 0.03 | 0.88 | 0.38 |
| Number of Adult Males Within 5m of Infant | -0.30 | 0.06 | -5.19 | **<0.0001** |
| Infant Age x Number of Adult Males | 0.03 | 0.01 | 2.68 | **0.007** |
| Maternal Dominance Rank | -0.07 | 0.07 | -1.00 | 0.32 |
| Maternal Age (Years) | 0.001 | 0.004 | 0.32 | 0.75 |
| ***Maternal Time Budget Controls (Reference = 100% feeding)*** |  |  |  |  |
| Prop. Time Mother Spent Resting | 0.39 | 0.06 | 6.47 | **<0.0001** |
| Prop. Time Mother Spent Walking | 0.24 | 0.06 | 4.25 | **<0.0001** |
| Prop. Time Mother Spent Socializing | 0.28 | 0.07 | 4.00 | **<0.0001** |

<sup>^</sup>Model also included random effects of mother ID with a random slope of infant age as well as a random effect of observer ID.
\*Observations were weighted by the number of minutes in each focal follow where it was known whether the infant was on the nipple.

**Table S2.** Predictors of the distance between an infant and its mother (square root transformed), estimated with a mixed effects model.

| **Parameter<sup>^</sup>** | **Coefficient Estimate** | **Std. Error** | **t value** | **p value** |
| --- | --- | --- | --- | --- |
| Intercept | -0.15 |  |  |  |
| ***Predictor Variables of interest*<sup>\*</sup>** |  |  |  |  |
| Infant Age (Months) | 0.36 | 0.02 | 19.6 | **<0.0001** |
| Infant Sex<br>(Reference = Female) | 0.22 | 0.09 | 2.53 | **0.01** |
| Number of Adult Males Within 5m of Infant | 0.58 | 0.08 | 6.85 | **<0.0001** |
| Maternal Age (Years) | 0.009 | 0.01 | 0.66 | 0.51 |
| Maternal Dominance Rank | 0.15 | 0.18 | 0.81 | 0.42 |
| ***Maternal Time Budget Controls (Reference = 100% feeding)<sup>\*</sup>*** |  |  |  |  |
| Prop. Time Mother Spent Resting | -1.08 | 0.17 | -6.40 | **<0.0001** |
| Prop. Time Mother Spent Walking | -1.27 | 0.16 | -7.88 | **<0.0001** |
| Prop. Time Mother Spent Socializing | -0.57 | 0.20 | -2.83 | **0.005** |

<sup>^</sup>Model also included random effects of mother ID and observer ID.
<sup>\*</sup>Observations were weighted by the number of minutes in each focal follow where the distance between mother and infant was known.

**Table S3.** Predictors of the Hinde Index of mother-infant proximity estimated with a mixed effects model.

| Parameter<sup>^</sup> | Coefficient Estimate | Std. Error | t value | p value |
| --- | --- | --- | --- | --- |
| **Hinde Index\*** |  |  |  |  |
| Intercept | -0.23 |  |  |  |
| ***Predictor Variables of interest*** |  |  |  |  |
| Infant Age (Months) | 0.06 | 0.007 | 8.53 | **<0.0001** |
| Infant Sex (Reference = Female) | 0.04 | 0.03 | 1.58 | 0.11 |
| Number of Adult Males Within 5m of Infant | -0.06 | 0.03 | -2.2 | **0.03** |
| Maternal Age (Years) | 0.0002 | 0.003 | 0.07 | 0.94 |
| Maternal Dominance Rank | 0.09 | 0.05 | 1.81 | 0.07 |
| ***Maternal Time Budget Controls (Reference = 100% feeding)*** |  |  |  |  |
| Prop. Time Mother Spent Resting | -0.11 | 0.05 | -2.05 | **0.04** |
| Prop. Time Mother Spent Walking | 0.08 | 0.05 | 1.47 | 0.14 |
| Prop. Time Mother Spent Socializing | -0.02 | 0.06 | -0.31 | 0.75 |

<sup>^</sup>Model also included random effects of mother ID along with a random slope of infant age as well as a random effect of observer ID.
\*Observations were weighted by the number of minutes in each focal follow where the distance between the mother and infant was known.

**Table S4.** Predictors of the time infants spent active, and of time spent feeding, walking, socializing and playing (the four components of the ‘active’ state; all arcsine square root transformed), estimated with mixed effects models.

| Parameter<sup>^</sup> | Response Variables- Infant Activity Budget<sup>*</sup> |  |  |  |  |
| --- | --- | --- | --- | --- | --- |
|  | Prop. of Time Infant Spent Active (Feeding + Walking + Socializing + Playing) | Prop. of Time Infant Spent Feeding | Prop. of Time Infant Spent Walking | Prop. of Time Infant Spent Socializing | Prop. of Time Infant Spent Playing |
| Intercept | 0.44 | 0.35 | 0.08 | 0.15 | 0.06 |
| **Predictor Variables of interest** |  |  |  |  |  |
| Infant Age (Months) | Coef. = 0.08<br>Std. Err = 0.006<br>t = 14.7<br>**p < 0.0001** | 0.06<br>0.004<br>14.3<br>**< 0.0001** | 0.04<br>0.004<br>11.4<br>**< 0.0001** | 0.002<br>0.003<br>0.46<br>0.65 | 0.02<br>0.003<br>6.17<br>**< 0.0001** |
| Infant Sex<br>(Reference = Female) | 0.007<br>0.03<br>0.28<br>0.78 | 0.006<br>0.02<br>0.40<br>0.69 | 0.02<br>0.02<br>1.26<br>0.21 | -0.005<br>0.015<br>-0.33<br>0.75 | 0.01<br>0.01<br>0.99<br>0.32 |
| Number of Adult Males Within<br>5m of Infant | 0.15<br>0.02<br>5.97<br>**< 0.0001** | 0.04<br>0.02<br>2.48<br>**0.01** | 0.06<br>0.02<br>3.78<br>**0.0002** | 0.07<br>0.02<br>4.47<br>**< 0.0001** | 0.07<br>0.01<br>4.49<br>**< 0.0001** |
| Maternal Age (Years) | 0.002<br>0.003<br>0.63<br>0.53 | 0.0006<br>0.002<br>0.31<br>0.76 | 0.002<br>0.002<br>0.78<br>0.44 | -0.004<br>0.002<br>-2.01<br>**0.04** | 0.003<br>0.002<br>1.63<br>0.10 |
| Maternal Dominance Rank | -0.004<br>0.05<br>-0.09<br>0.93 | 0.004<br>0.03<br>0.13<br>0.90 | 0.05<br>0.03<br>1.55<br>0.12 | -0.007<br>0.03<br>-0.25<br>0.80 | -0.02<br>0.02<br>-0.82<br>0.41 |
| **Maternal Time Budget Controls (Reference = 100% feeding)** |  |  |  |  |  |
| Prop. Time Mother Spent<br>Resting | -0.26<br>0.05<br>-5.17<br>**< 0.0001** | -0.41<br>0.04<br>-11.47<br>**< 0.0001** | -0.001<br>0.03<br>-0.04<br>0.97 | 0.20<br>0.03<br>6.45<br>**< 0.0001** | -0.09<br>0.03<br>-2.83<br>0.005 |
| Prop. Time Mother Spent<br>Walking | -0.49<br>0.05<br>-10.3<br>**< 0.0001** | -0.56<br>0.03<br>-16.0<br>**< 0.0001** | -0.04<br>0.03<br>-1.14<br>0.26 | 0.05<br>0.03<br>1.78<br>0.08 | -0.18<br>0.03<br>-6.21<br>**< 0.0001** |
| Prop. Time Mother Spent<br>Socializing | 0.07<br>0.06<br>1.18<br>0.24 | -0.33<br>0.04<br>-7.67<br>**< 0.0001** | 0.008<br>0.04<br>0.19<br>0.85 | 0.58<br>0.04<br>15.80<br>**< 0.0001** | 0.003<br>0.04<br>0.08<br>0.94 |

<sup>^</sup>Models also include random effects of mother ID with a random slope of infant age as well as a random effect of observer ID
<sup>\*</sup>Observations are weighted by the number of minutes in each focal follow where the infant’s activity was known.

**Table S5.** Predictors of the hourly frequency with which infants were engaged in positive and negative social interactions with non-mothers, estimated from two different mixed effects models.

| Parameter<sup>^</sup> | Coefficient Estimate | Std. Error | t value | p value |
| --- | --- | --- | --- | --- |
| **Positive Social Interactions with Non-Mothers Per Hour (square root transformed)\*** |  |  |  |  |
| Intercept | 3.3 |  |  |  |
| **Predictor Variables** |  |  |  |  |
| Infant Age (Months) | 0.5 | 0.1 | 4.15 | <0.0001 |
| Infant Age (Months) Squared | -0.06 | 0.01 | -4.64 | <0.0001 |
| Infant Sex (Reference = Female) | -0.2 | 0.2 | -0.33 | 0.75 |
| Number of Adult Males Within 5m of Infant | 1.3 | 0.2 | 6.94 | <0.0001 |
| Maternal Age (Years) | 0.007 | 0.03 | 0.28 | 0.78 |
| Maternal Dominance Rank | -0.4 | 0.4 | -1.0 | 0.31 |
| **Maternal Time Budget Controls (Reference = 100% feeding)** |  |  |  |  |
| Prop. Time Mother Spent Resting | 0.03 | 0.38 | 0.07 | 0.95 |
| Prop. Time Mother Spent Walking | -0.62 | 0.36 | -1.7 | 0.09 |
| Prop. Time Mother Spent Socializing | 3.04 | 0.45 | 6.76 | <0.0001 |
| **Negative Social Interactions with Non-Mothers Per Hour (not transformed)\*** |  |  |  |  |
| Intercept | 1.40 |  |  |  |
| **Predictor Variables** |  |  |  |  |
| Infant Age (Months) | -0.4 | 0.10 | -3.67 | 0.0003 |
| Infant Age (Months) Squared | 0.03 | 0.01 | 2.69 | 0.007 |
| Infant Sex (Reference = Female) | 0.04 | 0.13 | 0.32 | 0.75 |
| Number of Adult Males Within 5m of Infant | 0.68 | 0.15 | 4.54 | <0.0001 |
| Maternal Age (Years) | 0.002 | 0.02 | 0.12 | 0.91 |
| Maternal Dominance Rank | -0.49 | 0.24 | -1.97 | 0.046 |
| **Maternal Time Budget Controls (Reference = 100% feeding)** |  |  |  |  |
| Prop. Time Mother Spent Resting | 0.38 | 0.30 | 1.24 | 0.22 |
| Prop. Time Mother Spent Walking | 0.05 | 0.29 | 0.18 | 0.85 |
| Prop. Time Mother Spent Socializing | 2.08 | 0.36 | 5.71 | <0.0001 |

<sup>^</sup>Models also included random effects of mother ID and observer ID
\*Observations were weighted by the number of minutes in each focal follow where the infant's social behavior could be observed

**Table S6.** Predictors of the hourly frequency of social interactions between infants and adult males, and between infants and other non-mothers, estimated from two different mixed linear models.

| Parameter<sup>^</sup> | Coefficient Estimate | Std. Error | t value | p value |
| --- | --- | --- | --- | --- |
| **Social Interactions with Adult Males Per Hour (square root transformed)\*** |  |  |  |  |
| Intercept | 0.86 |  |  |  |
| **Predictor Variables** |  |  |  |  |
| Infant Age (Months) | -0.12 | 0.03 | -3.60 | **0.0003** |
| Infant Sex<br>(Reference = Female) | -0.15 | 0.16 | -0.96 | 0.34 |
| Number of Adult Males Within 5m of Infant | 3.22 | 0.24 | 13.61 | **<0.0001** |
| Maternal Age (Years) | -0.02 | 0.02 | -1.10 | 0.27 |
| Maternal Dominance Rank | 0.43 | 0.26 | 1.66 | 0.10 |
| **Maternal Time Budget Controls (Reference = 100% feeding)** |  |  |  |  |
| Prop. Time Mother Spent Resting | 0.58 | 0.42 | 1.40 | 0.16 |
| Prop. Time Mother Spent Walking | 0.46 | 0.33 | 1.39 | 0.16 |
| Prop. Time Mother Spent Socializing | 0.51 | 0.40 | 1.28 | 0.20 |
| **Social Interactions with Non-Mother Adult Females or Immatures Per Hour (square root transformed)\*** |  |  |  |  |
| Intercept | 5.52 |  |  |  |
| **Predictor Variables** |  |  |  |  |
| Infant Age (Months) Squared | -0.19 | 0.07 | -2.77 | **0.006** |
| Infant Sex<br>(Reference = Female) | 0.09 | 0.31 | 0.31 | 0.76 |
| Number of Adult Males Within 5m of Infant | 0.78 | 0.52 | 1.48 | 0.14 |
| Maternal Age (Years) | -0.0003 | 0.04 | -0.01 | 0.99 |
| Maternal Dominance Rank | -1.08 | 0.50 | -2.15 | **0.03** |
| **Maternal Time Budget Controls (Reference = 100% feeding)** |  |  |  |  |
| Prop. Time Mother Spent Resting | 0.78 | 0.90 | 0.86 | 0.39 |
| Prop. Time Mother Spent Walking | 0.22 | 0.72 | 0.30 | 0.76 |
| Prop. Time Mother Spent Socializing | 5.08 | 0.88 | 5.78 | **<0.0001** |

<sup>^</sup>Model also includes random effects of mother/infant pair ID (no duplicate mothers or multiple observers are contained in this subset of data).
\*Observations are weighted by the number of minutes in each focal follow where the infant's social behavior could be observed

**Table S7.**
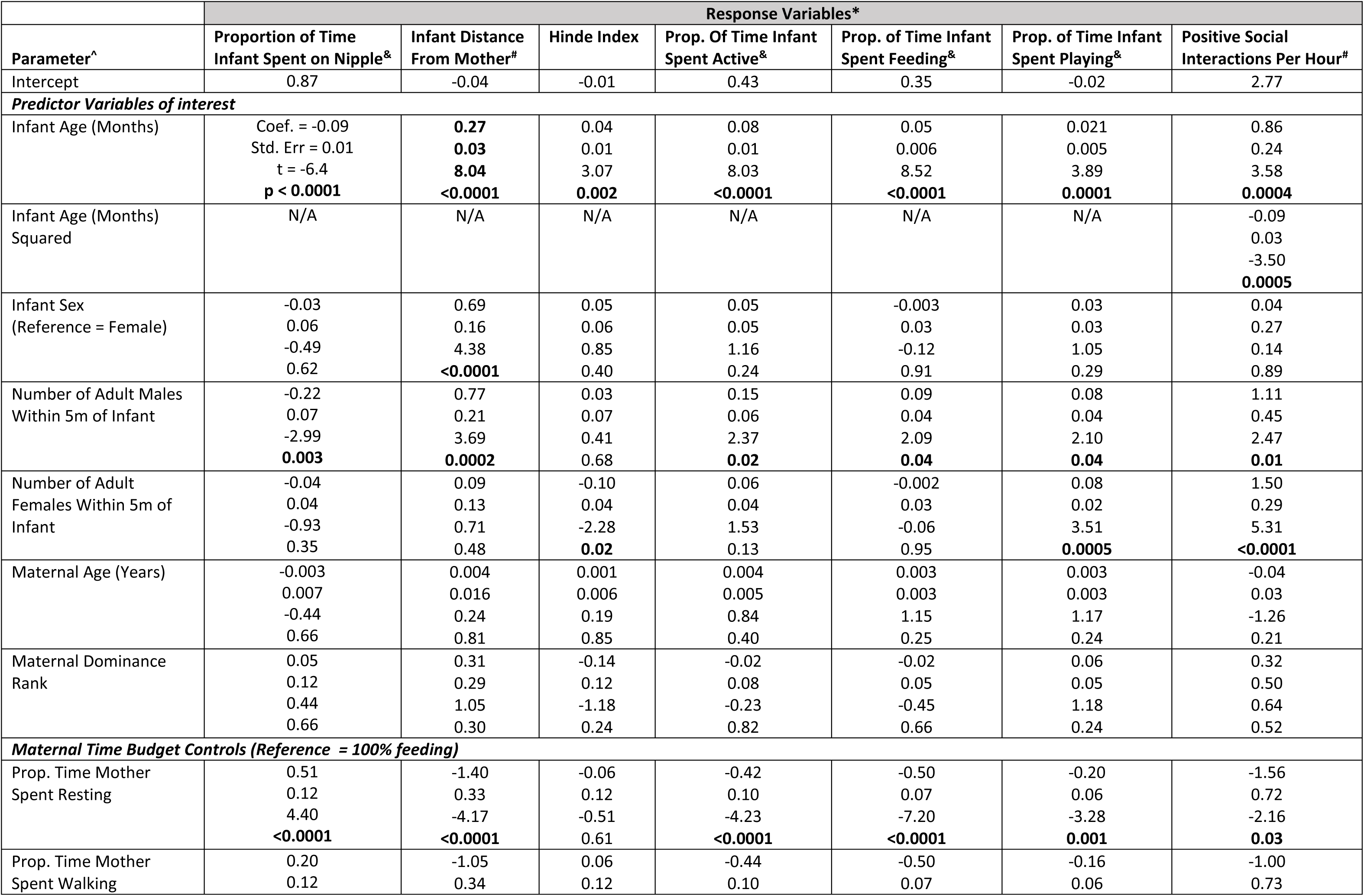
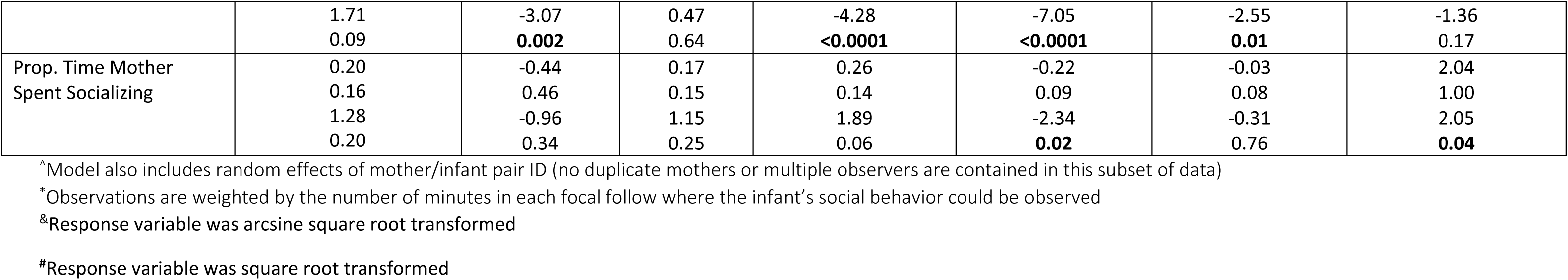
Results from mixed linear models of infant behavior that include both the number of adult males and adult females within 5m of the infant as predictor variables (available for 2021 data only, n = 289 focal follows).

**Table S8.**
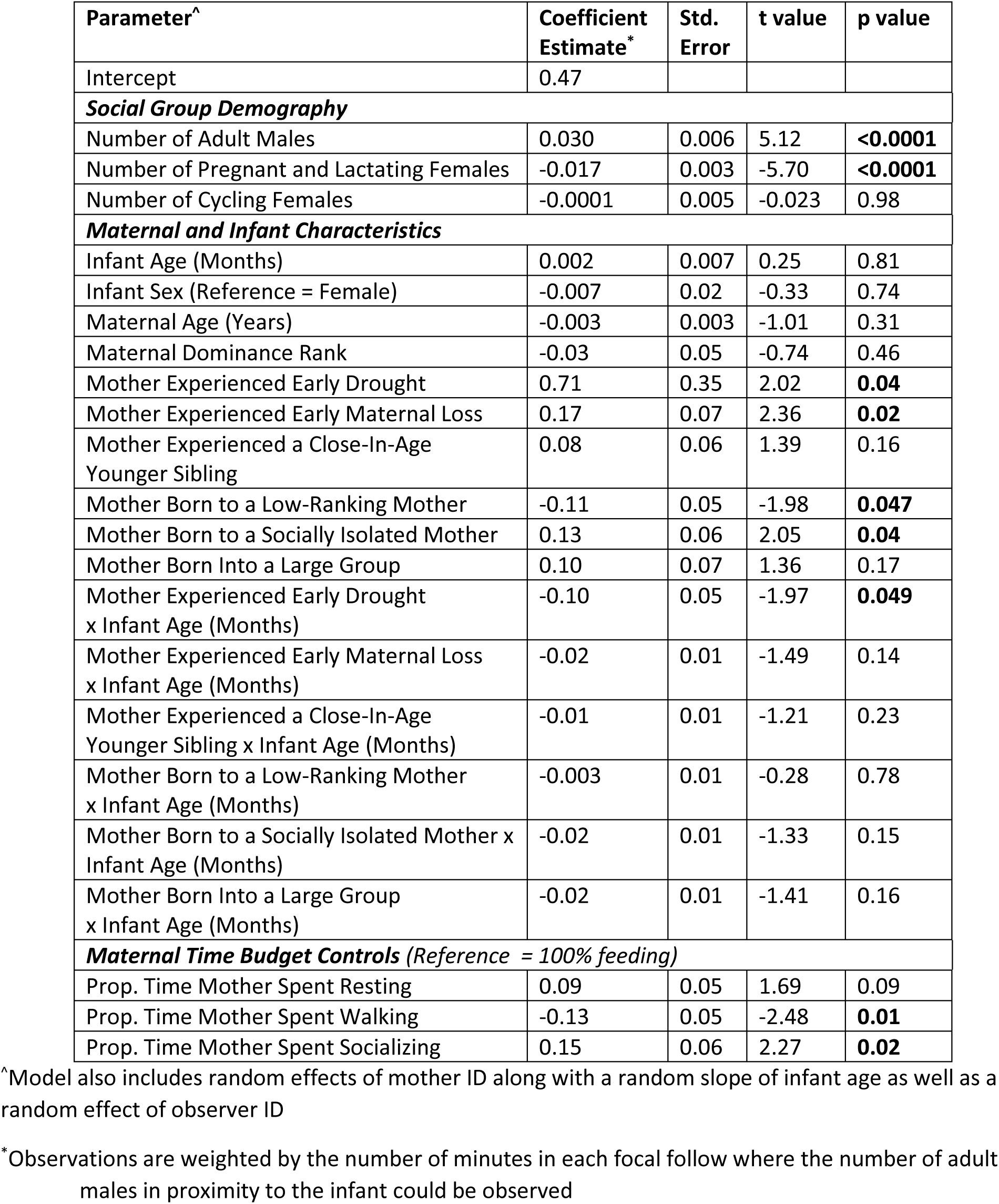
Predictors of the average number of males within 5 meters of an infant during a focal follow (square root transformed), with maternal early adversity expressed as 6 multivariate components, rather than a cumulative index.

### Supplementary Figures

**Figure S1.**
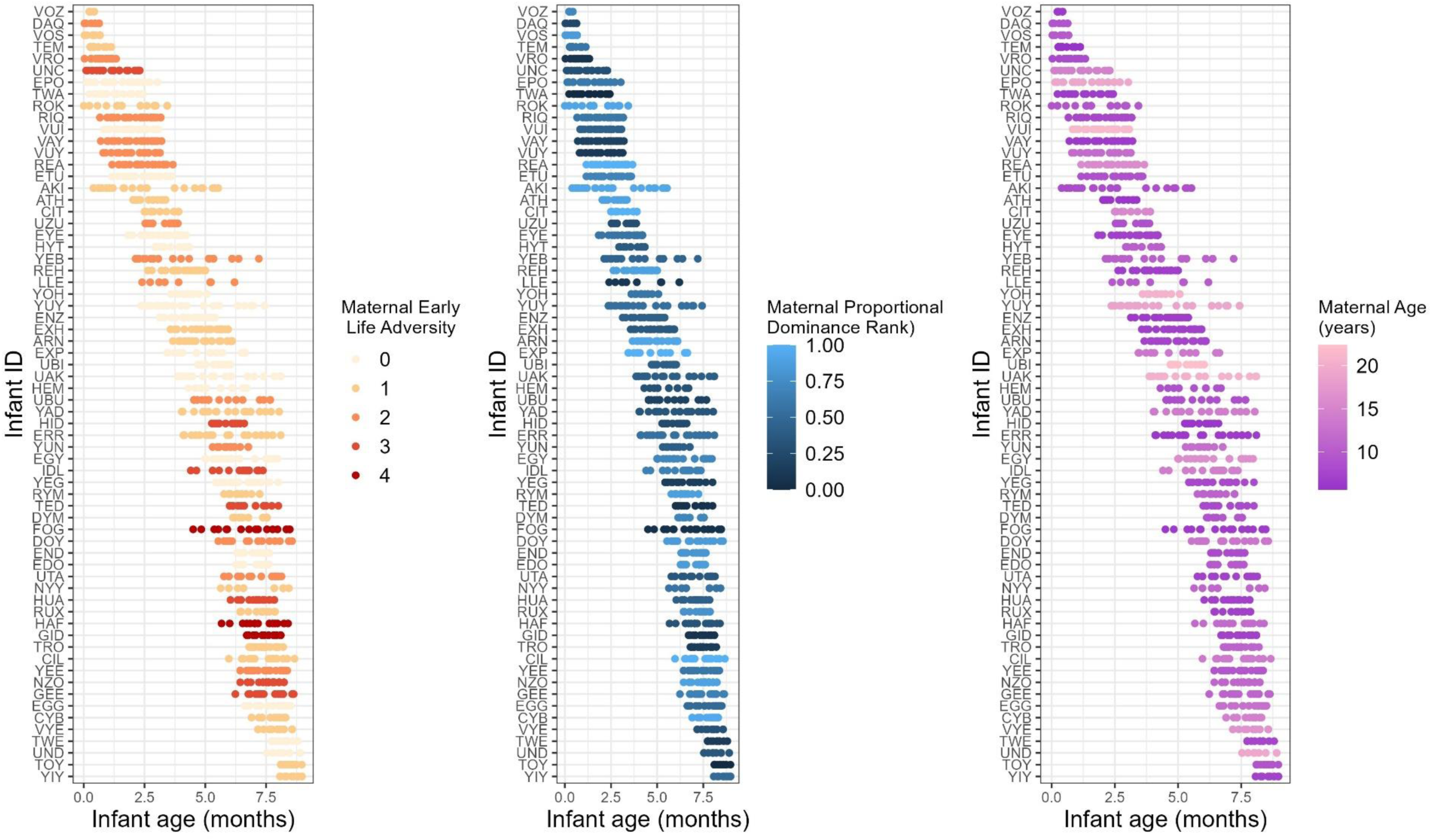
The behavioral dataset that supports all analyses in this paper. Each row represents an individual infant (n = 66), with each point (n = 1213) representing a focal follow. The shade of each dot represents either (left) the mother’s cumulative early adversity score, (middle) the mother’s proportional dominance rank, or (right) maternal age. The range of maternal early adversity measures, dominance ranks (0 = lowest ranking, 1 = highest ranking), and age were well represented across infants of different ages.

**Figure S2.**
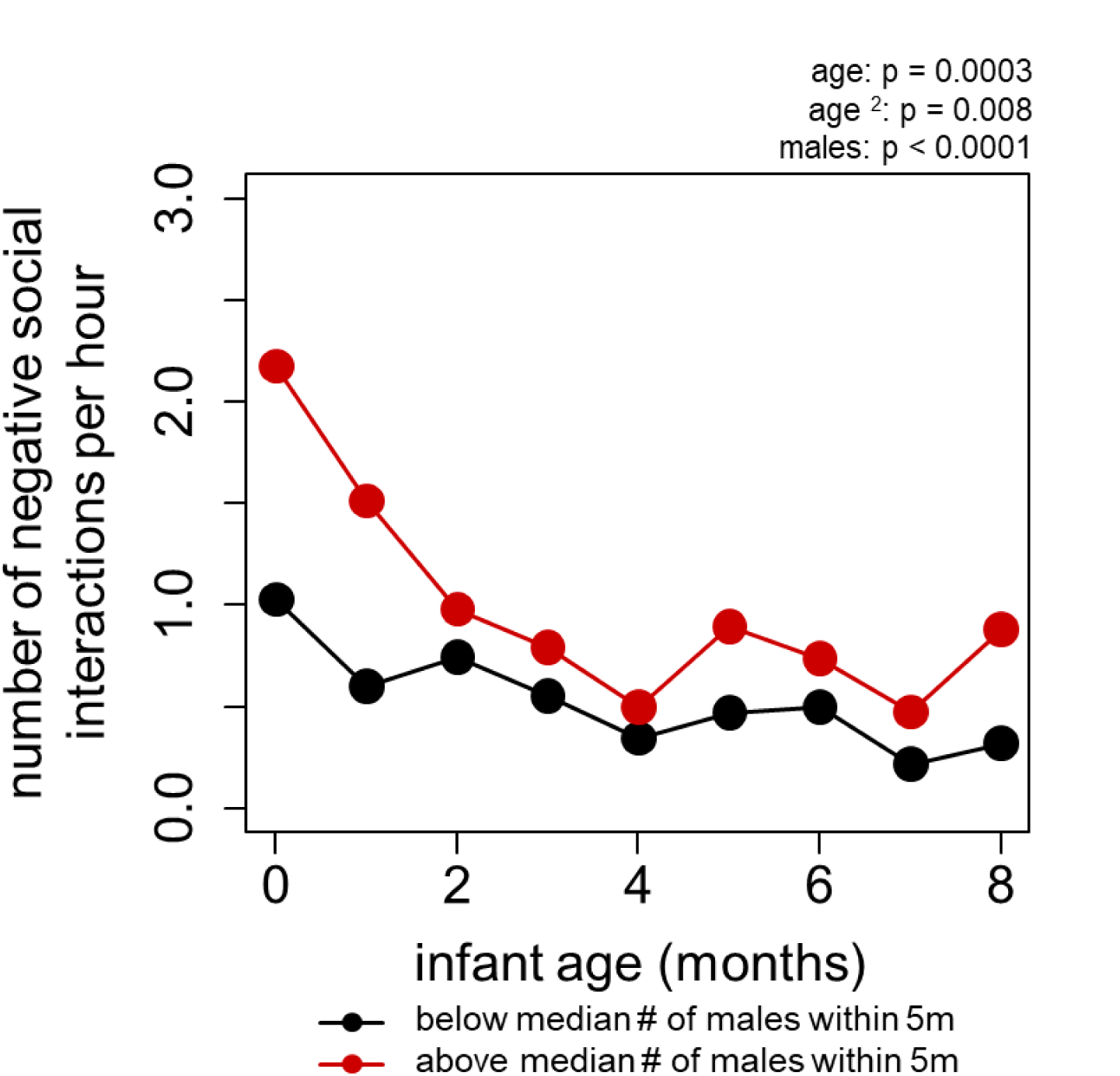
Infants engaged in more negative social interactions when they were in proximity to relatively many males. Points represent means across focal follows for infants of different ages. Each point represents infants that were contained in a one-month age window, which opens at the point’s x value (e.g. the points at x = 0 indicate infants that are between 0 and 1 months of age). p values indicate the results of the full models reported in Table S5.

